# All-night spectral and microstate EEG analysis in patients with Recurrent Isolated Sleep Paralysis (RISP)

**DOI:** 10.1101/2023.08.17.551215

**Authors:** Filip Černý, Václava Piorecká, Monika Kliková, Jana Kopřivová, Jitka Bušková, Marek Piorecký

## Abstract

The pathophysiology of recurrent isolated sleep paralysis (RISP) has yet to be fully clarified. Very little research has been performed on electroencephalographic (EEG) signatures outside RISP episodes. This study aimed to investigate whether sleep is disturbed even without the occurrence of a RISP episode and in a stage different than conventional REM sleep. 17 RISP patients and 17 control subjects underwent two consecutive full-night video-polysomnography recordings. Spectral analysis was performed on all sleep stages in the delta, theta, and alpha band. EEG microstate (MS) analysis was performed on the NREM 3 phase due to the overall high correlation of subject template maps with canonical templates. Spectral analysis showed a significantly higher power of theta band activity in REM and NREM 2 sleep stages in RISP patients. The observed rise was also apparent in other sleep stages. Conversely, alpha power showed a downward trend in RISP patients’ deep sleep. MS maps similar to canonical topographies were obtained indicating the preservation of prototypical EEG generators in RISP patients. RISP patients showed significant differences in the temporal dynamics of MS, expressed by different transitions between MS C and D and between MS A and B. Both spectral analysis and MS characteristics showed abnormalities in the sleep of non-episodic RISP subjects. Our findings suggest that in order to understand the neurobiological background of RISP, there is a need to extend the analyses beyond REM-related processes and highlight the value of EEG microstate dynamics as promising functional biomarkers of RISP.

**Significance Statement:** We focused on tracking electrophysiological traces of RISP (a REM parasomnia) beyond REM sleep of subjects clinically diagnosed with RISP outside of RISP episodes. We observed a rise of theta band activity in NREM 2 sleep of RISP patients. This may point to a larger dysregulation of sleep mechanism making the person more prone to sudden awakenings in the upcoming REM sleep. Theta band differences were further observed in REM sleep. We additionally utilized the EEG MS methodology on deep sleep to investigate differences in dominant brain topographies. Though dominant brain topographies are consistent with canonical MS, RISP patients show significantly different transitioning between sleep-related topographies suggesting a difference in their sleep regulation mechanisms.

## Introduction

Recurrent isolated sleep paralysis (RISP) is a sleep disorder whose mechanism has not yet been fully elucidated due to its random occurrence and difficulty to capture in laboratory conditions [1]. It manifests by repeated episodes with an inability to perform voluntary movements at sleep onset or on a sudden awakening from sleep. It is often accompanied by hallucinatory experiences, e.g., feeling someone’s presence, or feeling pressure against the chest [2]. Although RISP is not physically harmful, it can leave the victim with anxiety and stress [3]. Research shows that the experience of an isolated sleep paralysis episode is about 8 % [4] in the general population. However, it was shown to be higher in psychiatric patients [5]. A frequent occurrence in the population [5, 6] and the association with clinically significant distress make RISP an important research topic [7]. However, very few studies have addressed the electrophysiological correlates of RISP episodes or even examined brain activity outside of them.

The RISP episodes have primarily been described as dissociative states of consciousness, combining a mixture of waking and REM sleep brain states with abundant alpha activity in electroencephalography (EEG) and persistence of muscle atonia [8, 9]. Although recently, a predominant theta has been detected during the actual episodes using spectral analysis [10], which may indicate that the brain is more likely to be in dreaming than waking state. [8] concluded a case study on a narcolepsy patient, suggesting that sleep paralysis represents a transition state between REM sleep and wakefulness. A study by [9] shows a possible higher paralysis probability in sudden interruptions of REM sleep.

Patients suffering from sleep apnea also seem to be more prone to sleep paralysis [11]. This may have been due to higher REM sleep fragmentation as apneic breaks occur during REM sleep 36 % of the time [12]. However, higher REM sleep fragmentation was not present in our previous study of RISP patients [13] in REM sleep along with differences in other macrostructural sleep parameters. Nevertheless, we observed differences in EEG spectral bands between RISP patients and control subjects. Patients showed higher bifrontal beta activity in the patients’ group in REM sleep. Persistent higher cortical activity may represent a predisposing factor for sleep paralysis [13].

On the other hand, sleep paralysis may not be a matter of REM sleep alone and may reflect global sleep dysregulation. Its incidence is known to increase with elevated homeostatic pressure, i.e., due to sleep deprivation, irregular sleep-wake schedules, and jet lag. Avoiding these predisposing factors is also the most effective therapy for RISP patients [14]. RISP also often occurs as a symptom of narcolepsy, a disease characterized by disruption of the circadian sleep-wake rhythm [15] or interruption of the REM-NREM cycle that may favor sleep paralysis. Changes in EEG NREM sleep oscillations were also found during nightmares [16, 17], which may be a decisive predictive factor for the frequency of RISP episodes [18].

Our aim was to determine whether differences in the global functional state of the sleeping brain may play a role in the pathophysiology of RISP regardless of the presence of episodes or even sleep stages and to describe their dynamics. In contrast to our previous work on this dataset [13], focusing on the REM stage, here we asked whether this REM sleep parasomnia also shows differences in NREM sleep which may suggest a broader, more complex dysregulation of sleep than previously thought.

## Materials and Methods

### Experimental Design

All study participants were invited through advertisements on the website of the National Institute of Mental Health website, Klecany, and social media networks. Subjects with a history of sleep paralysis episodes (N = 112) underwent a clinical interview that served as a diagnostic tool to verify that participants fulfilled the criteria for RISP [4]. A total of 20 selected participants with RISP underwent two consecutive full-night video-polysomnography (vPSG) with consecutive multiple sleep latency tests (MSLT) to exclude narcolepsy. Inclusion criteria contained fulfillment of RISP standards defined in the International Classification of Sleep Disorders 3 (ICSD-3) [4] and experience of 2 sleep paralysis (SP) episodes in the past six months as recommended by [7]. Exclusion criteria contained a diagnosis of other sleep disorders, clinically significant neurological, psychiatric, or somatic disorders that could lead to sleep alteration, and medication affecting sleep (e.g., antidepressants and anxiolytics). The control group comprised 20 healthy age-matched volunteers with no history of SP episodes; who did not experience neurological, psychiatric, or sleep disorders; and who did not receive any psychotropic medication. Before analysis, we excluded one participant from the RISP group due to consuming a benzodiazepine drug before the experimental night and one participant from the control group due to technical issues. Furthermore, we excluded two participants from each group due to uncorrelated MS topographical maps with templates (see Microstate validation).

In total, 17 participants in the RISP group (N = 15 women, 2 men; mean age = 25.24 years, SD = 7.0) and 17 healthy participants in the control group (N= 13 women, 4 men; mean age = 25.17, SD = 6.20) were included. The frequencies of SP episodes were several times a year (N = 5), once a month (N = 5), and several times a month (N = 7). Six months before the PSG recording, participants experienced a mean of 10.07 SP episodes (SD 8.04) ranging from 2 to 30 episodes (one outlier stated experience of 70 episodes). There was no between-group difference in gender and age.

The first night of vPSG served as an adaptation night and for the exclusion of other sleep disorders. Therefore, only data from the second night (experimental) were analyzed. VPSG was performed using a digital Brainscope PSG system (M&I spol. s.r.o., Czech Republic). It consisted of 19-channel EEG, electrooculography, electromyography of the bilateral mentalis muscle, the bilateral flexor digitorum superficialis muscle, and the bilateral tibialis anterior muscle, electrocardiography, airflow, thoracic and abdominal respiratory effort, oxygen saturation, microphone and synchronized video monitoring from 10:00 pm to 06:00 am, according to the recommendation of the American Academy of Sleep Medicine (AASM) [4]. MSLT measurements were taken in agreement with standard protocol at 09:00 am, 11:00 am, 01:00 pm, 03:00 pm, and 05:00 pm.

After adaptation night, participants were allowed to leave the sleep laboratory in the morning and return at 06:00 pm. They were encouraged to follow their usual daily activities and sleep hygiene rules.

The study was conducted in accordance with the Declaration of Helsinki. All participants provided written informed consent. The study protocol was approved by the local Ethical Committee of the National Institute of Mental Health, Klecany, Czech Republic, and by the Ethical Committee of the Third Faculty of Medicine, Charles University in Prague, Czech Republic. A financial reward of 1500 CZK (approximately 78 $) was provided for participation in polysomnographic recordings.

### Data preprocessing

EEG data were recorded at a sample rate of 1000 Hz and downsampled to 250 Hz. Slow drifts of the signal, as well as line noise and other high-frequency noises, were filtered out by a 0.5-30 Hz bandpass FIR filter with an order of 1000 in the context of spectral analysis and 1-40 Hz in the context of microstate analysis and literature evaluation [19, 20, 21]. Each polysomnographic recording was scored for ongoing sleep phases by a somnologist, and each epoch was assigned into groups corresponding to the ongoing sleep phase: NREM 1, NREM 2, NREM 3, REM, and WAKE based on the sleep scoring results.

### EEG Spectral Analysis

For spectral analysis, EEG signals were evaluated in a montage to the mastoid electrodes (M1, M2). Data were filtered with a two-way bandpass type FIR filter in the 0.5-30.0 Hz frequency range.

EEG spectra were evaluated using power spectral density analysis. Power spectra were estimated by frequency analysis using conventional multiple windows based on discrete prolate spheroidal sequences or Slepian sequences (DPSS) [22, 23]. Selected frequency bands of interest included delta (0.5-4.0 Hz), theta (4.0-8.0 Hz), and alpha (8.0-12.0 Hz) EEG bands, which cover the majority of sleep-related activity. Beta band was not analyzed due to previously shown indifferences in between RISP and sleep [10]. Power spectral densities were further converted to relative power spectra. Relative spectrum represents a percentage of power in a frequency band and enables precise statistical comparison of activity between bands [24].

### EEG microstates analysis

As a complementary approach to spectral analysis, we performed EEG microstates (MS) analysis. EEG MS represent short periods of quasi-stability of electric field configurations that reflect a reference-free global functional state of the brain. They are thought to arise due to the coordinated activity of neuronal assemblies originating from large cortical areas and have distinct scalp topographies replicated by numerous studies [25]. Temporal dynamics of EEG MS are altered in various states of consciousness [20]. If the brain activity of RISP patients differs during sleep, even outside the paralysis episode, microstates could reflect these differences. Even though RISP is mainly associated with the REM phase, previous studies analyzed the REM and NREM together [26, 27, 28]. Furthermore, NREM 3 sleep phase has been associated with REM phase analysis in depression research [29] and in a sleep quality study [30].

We performed the analysis only for the NREM 3 stage based on template maps correlations (see Microstate validation chapter). The average electrode reference was used, and data were bandpass filtered in a range of 1-40 Hz [19, 20, 21]. Demean and detrend filters were used as well. The FieldTrip toolbox [22] was used for preprocessing purposes. The final control’s group mean NREM 3 record duration was 94.90 min (SD 21.77), totaling 1613.22 min. RISP patients’ group mean NREM 3 record duration was 88.36 min (SD 23.11), totaling 1502.20 min.

The modified k-means algorithm was used to cluster the resulting topographical maps in peaks of the global field power (GFP) curve [31]. This algorithm was restarted 20 times. The polarity of EEG curves was ignored. Standard EEG microstate parameters (duration, occurrence, coverage, GFP, transitions) [32] were evaluated for each microstate class.

### Microstate validation

To further support the EEG microstate approach, we tested the degree to which the individual’s activity corresponded to the created microstate templates. In other words, we verified whether stable results were obtained for all MS topographies. Individual subject topographies were therefore compared with group template maps (RISP group and control maps separately) and with Grand mean (GM) global maps of the whole dataset. To do this, we used spatial correlation analysis and visual comparison. The spatial correlation was computed using the global dissimilarity index (GDI) [33], defined as:

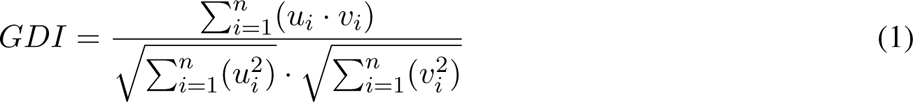

where *u* and *v* represent the measured voltage under electrodes, *i* represents individual electrode and *n* represents the total number of electrodes. This parameter corresponds to the spatial Pearson’s product-moment correlation coefficient [20].

### Statistical analysis

Considering that there are no studies that present robust sleep EEG datasets of RISP patients, we performed power analysis on other sleep disorders. Insomnia, which has various spectral characteristics across different EEG bands described, was selected. We hypothesized to observe a difference in the theta band, which is typical for the REM phase. We evaluated the relative spectral performances for 4 sleep stages (REM, NREM1, NREM2, and NREM3) and 3 EEG bands (delta, theta, and alpha), due to relative spectral power being a more sensitive marker for NREM sleep [34]. Studies included in the review of [34] use various ranges of spectral bands, therefore the calculation of the sample size was done through the combined mean values and standard deviations. Effect sizes were determined based on Hedges’G [35]. Power analysis estimates for relative spectra were transitioned for MS analysis, as the area of interest was based on spectral analysis results.

This review suggests a group of 30 insomniacs and 20 healthy controls to be necessary to identify a difference in the REM band for relative theta spectral power. As RISP is not as widespread as insomnia it is challenging to select age and gender-matched patients who only suffer from this disease without other neuropsychiatric disorders. We, therefore, chose a dataset size of 20 controls and 20 patients. Due to technical and data acquisition issues, we excluded a few subjects. In total, 17 patients and 17 controls entered the analysis.

Results from EEG spectral analysis were statistically evaluated on two levels. For the normalized EEG power spectra, we performed permutation tests of the FieldTrip toolbox (Monte-Carlo estimates of the probability of significance and critical values) with cluster correction [22, 36]. A T value was computed for every channel), clustered based on spatial adjacency, and compared between experimental conditions to evaluate its effect. Secondly, false discovery rate (FDR) correction was used to correct for multiple comparisons of spectral analysis results between different spectral ranges and sleep phases. Thus we performed corrections for multiple channels, spectra, and sleep phases.

Temporal parameters that describe EEG MS maps were statistically evaluated between groups. The non-parametric Wilcoxon rank sum test for two populations, where samples are independent, was used. The resulting p-values were corrected for multiple comparisons with Bonferroni’s correction. The correction coefficient equaled 16 for temporal parameters of occurrence, duration, contribution, and GFP (4 MS x 4 parameters). Transition parameters were corrected by the coefficient equal to 16 following both directions of transitions without autocorrelations [37].

### Code accessibility

All processing scripts are available on: https://github.com/vaclavapiorecka/sleepo.

## Results

### Macrostructural vPSG sleep parameters

During the experimental nights, no episodes corresponding to RISP diagnosis were detected. MSLT results excluded narcolepsy in all RISP participants. Table 1 shows the mean values of macrostructural sleep parameters.

**Table 1:** Macrostructural sleep parameters of RISP (n=17) and control group (n=17). Displayed are mean and standard deviations. SOL-sleep onset latency, TST-total sleep time. There were no statistically significant differences between the groups.

|  | RISP patients | Controls | p-value |
| --- | --- | --- | --- |
| **SOL (min)** | 22.93±13.83 | 21.43±19.61 | 0.69 |
| **TST (min)** | 422.59±23.01 | 422.29±38.01 | 0.98 |
| **Sleep efficiency (%)** | 94.76±2.43 | 93.56±4.02 | 0.30 |
| **REM (%)** | 21.09±3.90 | 22.22±3.69 | 0.39 |
| **N1 (%)** | 4.15±2.27 | 3.29±1.35 | 0.19 |
| **N1 LAT (min)** | 8.71±19.93 | 21.62±44.15 | 0.28 |
| **N2 (%)** | 44.45±7.77 | 40.68±7.41 | 0.16 |
| **N2 LAT (min)** | 2.00±1.87 | 2.12±2.23 | 0.87 |
| **N3 (%)** | 25.05±3.98 | 27.34±4.60 | 0.13 |
| **N3 LAT (min)** | 14.16±4.02 | 11.21±3.41 | 0.03 |
| **Wake (%)** | 5.24±2.43 | 6.44±4.02 | 0.30 |
| **Wake LAT (min)** | 24.12±25.02 | 43.88±34.87 | 0.07 |

### EEG spectral analysis

Power values of EEG trials were computed for three defined spectral bands of interest. Spectral power was calculated for trials in each sleep stage individually. Figure 1 shows a topographically mapped difference in spectral power between the RISP group and controls. There were no statistically significant differences when using only the Bonferroni correction (19 electrodes x 4 sleep stages x 3 spectral bands); this is, however, a very strict correction. Cluster-based and FDR-corrected results showed significantly higher activity in the theta band for the NREM 2 and REM phases. A rise in theta band activity was apparent across the whole scalp for the NREM 2 phase.

**Figure 1:**
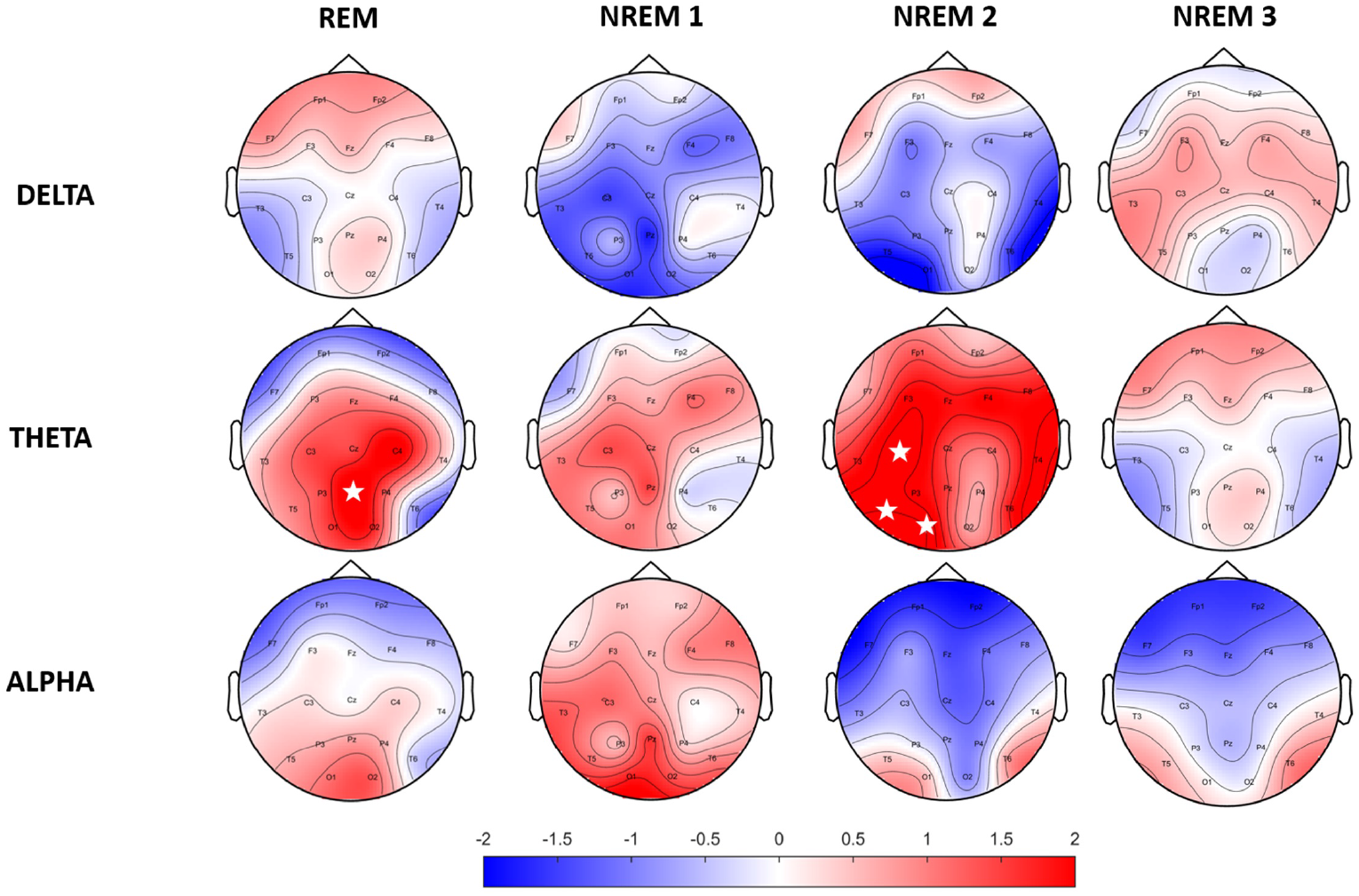
Topographic maps of relative spectral power differences between RISP patients group and healthy controls. Positive values represent higher activity in the RISP patients group. White asterisks mark electrodes with statistically significant differences after FDR correction.

### EEG microstate analysis

Identified MS maps represented 80,29 ± 1,56 % GEV for the patients’ group and 79,00 ± 1,85 % for the control group in the case of four template maps. Based on literature [38], GEV, and visual inspection of MS templates, this number of maps was chosen as sufficient for analysis. Therefore, this number of template maps was extracted for both groups (controls and RISP patients). The topographies of both groups of microstates and GM topographies are depicted in Figure 2 and compared to canonical MS maps of [38] in Figure 3.

**Figure 2:**
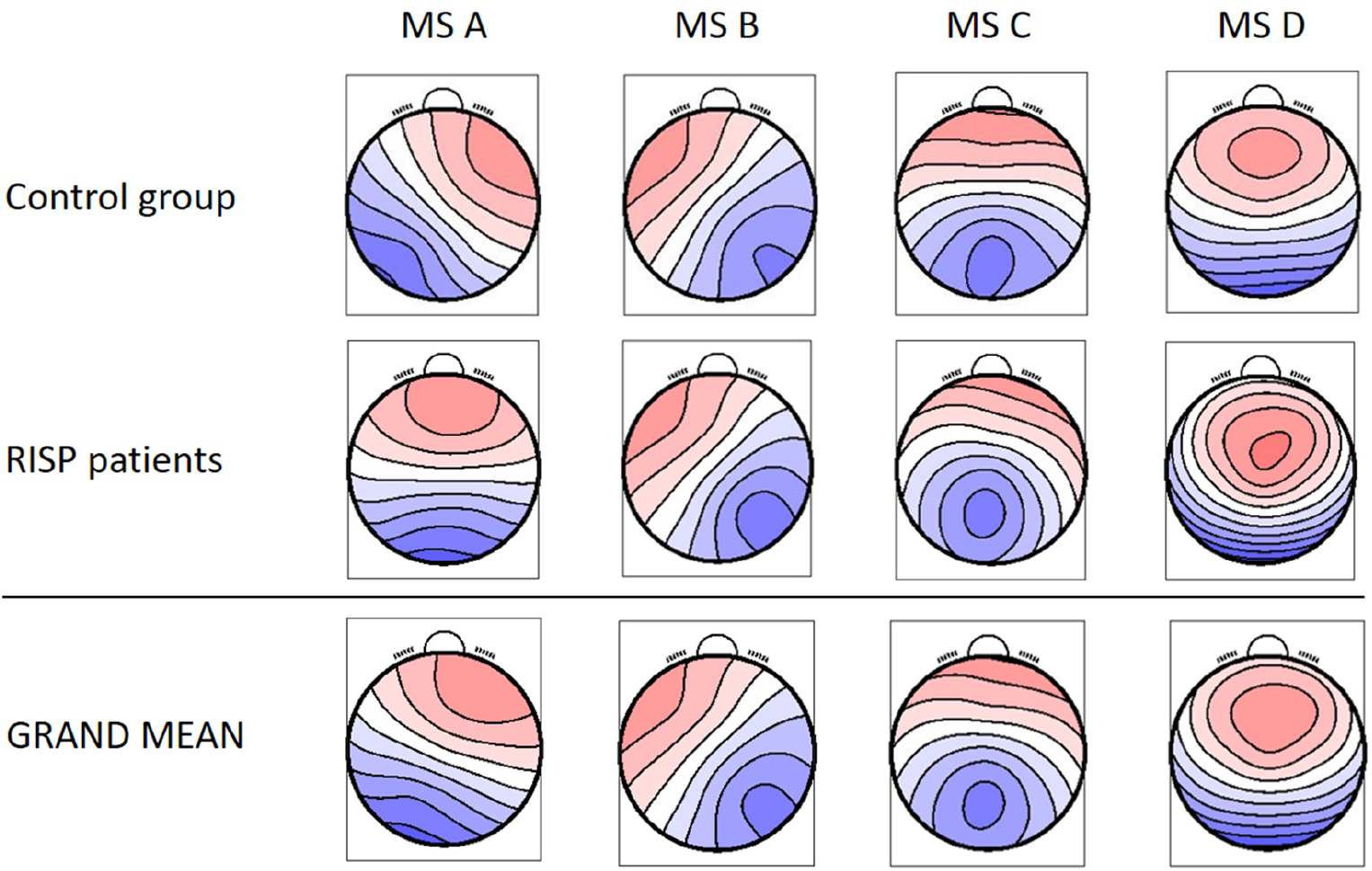
Resulting four MS topographical maps for individual groups and the GM MS maps for the whole dataset. Created with EEG microstates plugin for EEGlab.

**Figure 3:**
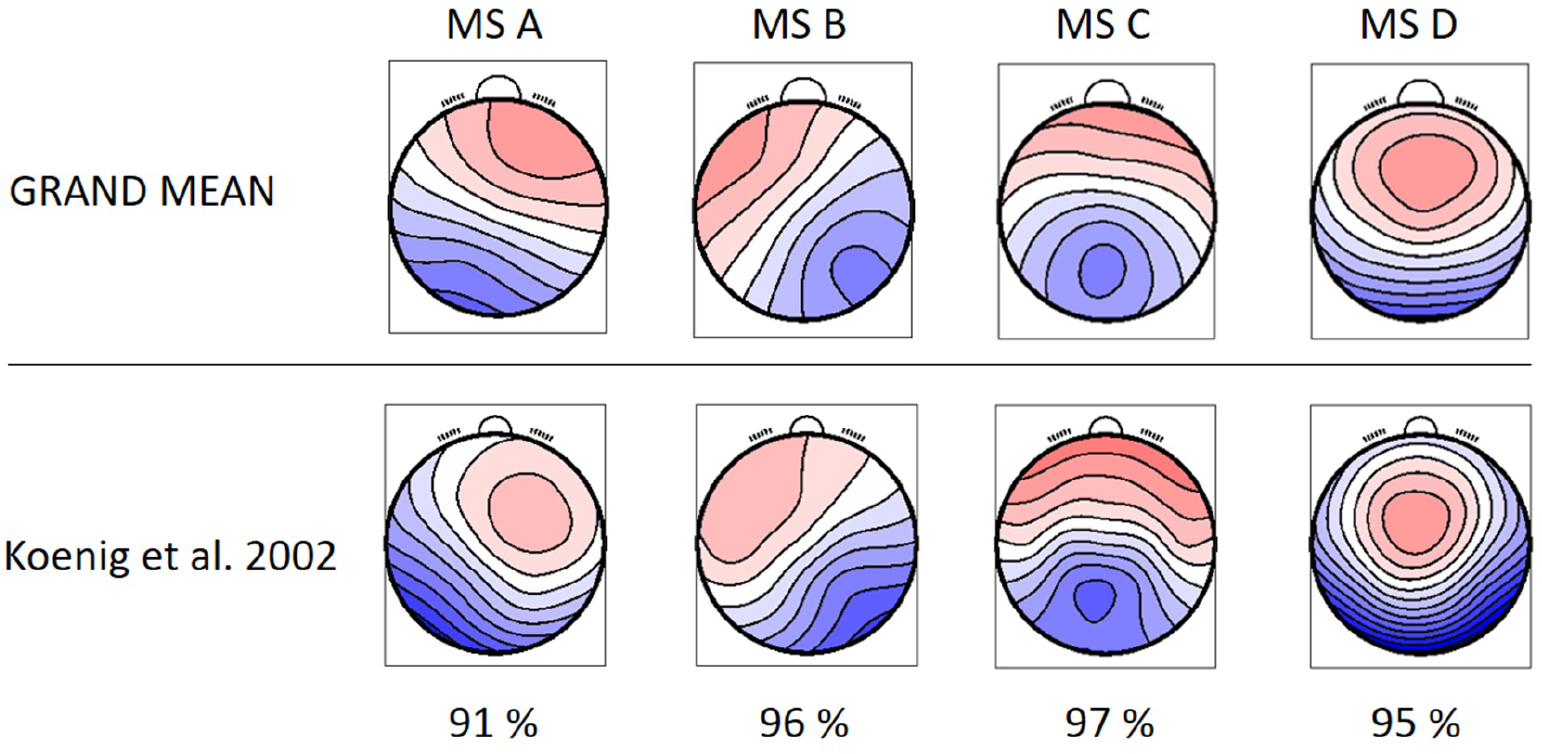
GM topographical maps of RISP and healthy controls in comparison to the topographical maps extracted by [38]. The percentage of spatial correlation for each microstate topography is shown below.

The resulting MS, derived as an average topography of both groups, have similar topographies to those standardized in the literature (A-D) [38, 39, 40, 41] and thus were labeled accordingly. The spatial correlation of the GM maps with the topographies of individual groups is presented in Figure 4.

**Figure 4:**
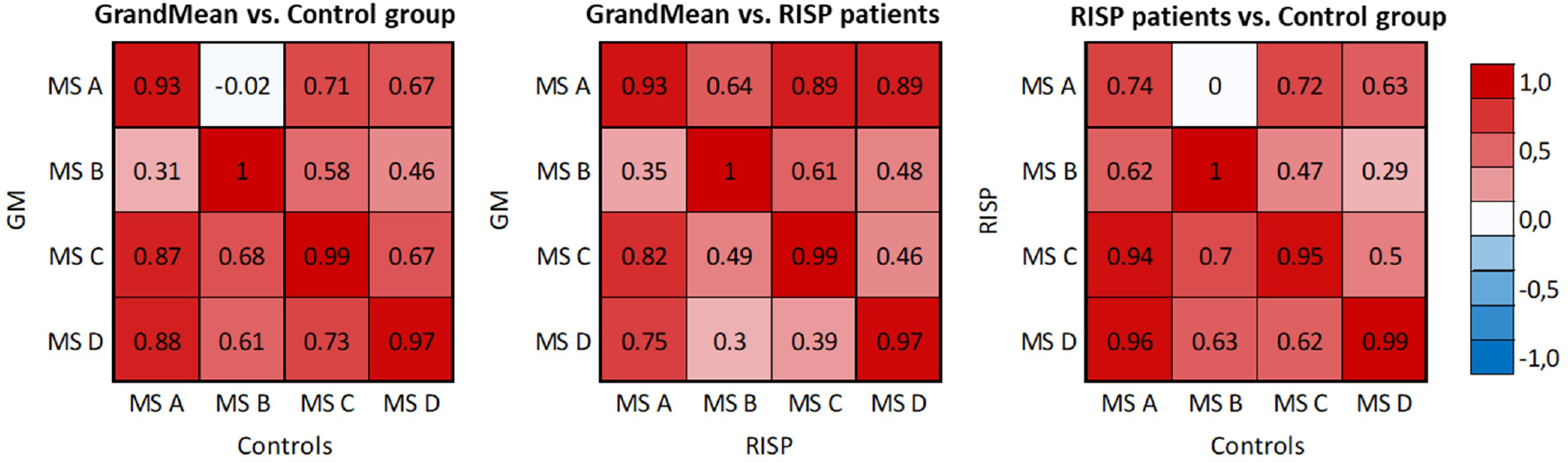
Spatial correlation matrices of identified MS templates of RISP patients, control group, and GM of the whole dataset. The scale was set between −1 and 1.

The correlation matrices describe correlations between the microstate topography of the RISP patients and control group and the correlation of each group and the GM microstate topographies. The correlation of both groups’ topographies and GM topographies validated that using the GM template for sorting purposes is adequate.

**Table 2:** MS temporal parameters for RISP patients (upper part of the row) and control group. GEV for RISP patients was 80.3±1.56 % and 79.0±1.85 % for control group.

|  | MS A | MS B | MS C | MS D |
| --- | --- | --- | --- | --- |
| **Coverage (%)** | $0.24 \pm 0.02$ | $0.25 \pm 0.03$ | $0.27 \pm 0.04$ | $0.25 \pm 0.04$ |
| | $0.21 \pm 0.03$ | $0.24 \pm 0.02$ | $0.27 \pm 0.03$ | $0.27 \pm 0.02$ |
| **Duration (ms)** | $88.7 \pm 9.34$ | $87.7 \pm 10.4$ | $92.2 \pm 12.4$ | $87.6 \pm 9.19$ |
| | $86.8 \pm 8.88$ | $87.1 \pm 8.39$ | $94.7 \pm 9.81$ | $93.0 \pm 9.18$ |
| **Occurrence ($s^{-1}$)** | $2.72 \pm 0.31$ | $2.96 \pm 0.32$ | $2.96 \pm 0.33$ | $2.86 \pm 0.42$ |
| | $2.52 \pm 0.30$ | $2.82 \pm 0.26$ | $2.98 \pm 0.28$ | $3.03 \pm 0.29$ |
| **GFP ($\mu V$)** | $10.4 \pm 2.43$ | $9.48 \pm 2.08$ | $10.4 \pm 2.62$ | $10.0 \pm 2.14$ |
| | $10.79 \pm 2.81$ | $9.86 \pm 2.49$ | $10.8 \pm 2.80$ | $10.7 \pm 2.70$ |

The mean and SD values of MS parameters duration, occurrence, coverage, and GFP were computed across the groups, see Table 2. The mean values of transitions are visualized in Figure 5. Due to the non-normal probability distribution, the non-parametric test for two independent samples was used, specifically the Wilcoxon rank sum test. The resulting p-values are visualized in Table 3 and Figure 5.

Spatial correlations were computed for each MS in single-group analyses. This way we checked for outlier maps, such as maps from data with artifacts. MS A and B are uncorrelated in the case of normative templates [38] (0.04) and also in our control group (−0.04). At the same time, there is already a weak spatial correlation (0.66) in the RISP group. In RISP, the transition between A-B is reduced. Controls have a higher spatial correlation between MS C and D (0.82), while in RISP subjects MS C and D are uncorrelated (0.28). The C-D transition is elevated in RISP. The transition probability between MS C and D in RISP was negatively related to the degree of the difference in correlation between controls and RISP. When the correlation between these MS is lower, their transitions are more frequent.

**Table 3:** Statistical evaluation of MS parameters between controls and RISP group. Displayed are the p-values of the Wilcoxon rank sum test. The p-value of 0.05 is corrected for multiple comparisons to 0.004167.s.

|  | MS A | MS B | MS C | MS D |
| --- | --- | --- | --- | --- |
| **Coverage** | 0.03 | 0.19 | 0.45 | 0.02 |
| **Duration** | 0.54 | 0.84 | 0.43 | 0.19 |
| **Occurence** | 0.07 | 0.24 | 0.63 | 0.16 |
| **GFP** | 0.63 | 0.63 | 0.49 | 0.39 |

**Figure 5:**
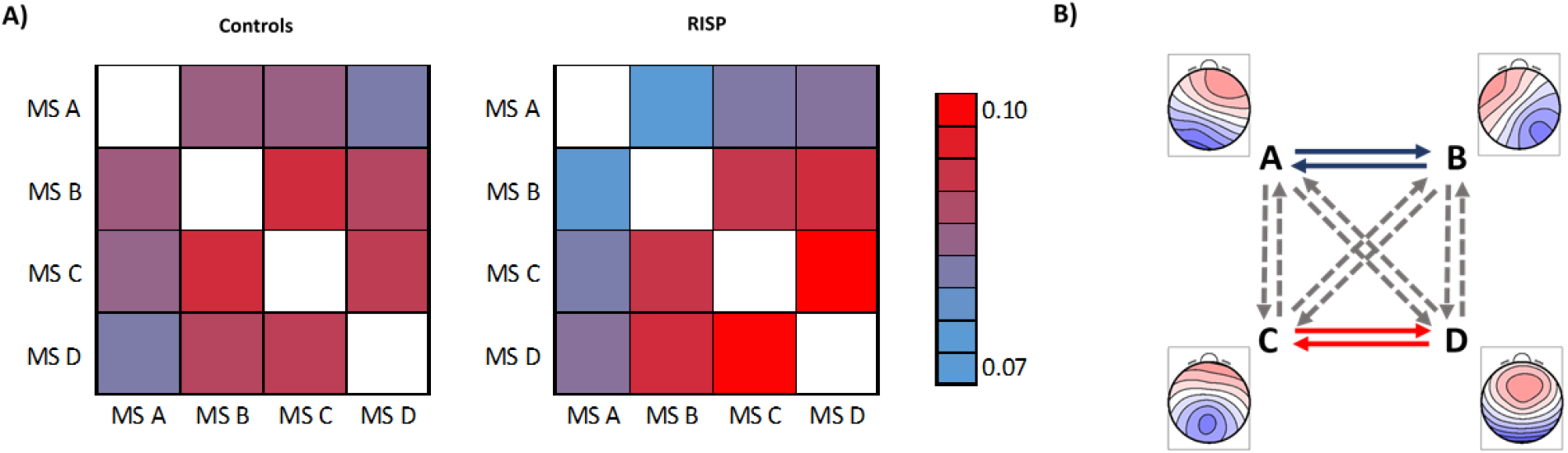
Mean transitions between MS. Part A) left represents mean transitions of control and A) right RISP group. B) Statistically significant differences of transition parameters between healthy controls and RISP patients group, represented by GM microstates topographical maps. The p-value of 0.05 is corrected for the multiple comparisons to 0.0042.

In broadband, we observed significant differences in the transition between MS A and B and between MS C and D in both directions. The patient group showed a lower transition rate between MS A and B and a higher transition rate between MS C and D than the control group.

## Discussion

In this study, we present an analysis of EEG sleep parameters of patients diagnosed with RISP. In current literature [6, 42], RISP has mostly been associated with REM sleep, however, the precise mechanism underlying this disorder is unknown. Compared to other studies [6, 13] focusing primarily on REM sleep, we targeted the NREM sleep stages using multiple analysis methods on a unique dataset obtained with strict entry criteria.

EEG spectral analysis showed a significantly higher theta activity in REM and NREM 2 sleep stages in RISP patients compared to healthy controls. This finding corresponds with the results of [10] describing predominant theta activity during a paralysis episode [43]. Our patients did not, however, experience any paralysis episodes during the recording night. A higher occurrence of theta activity over the scalp compared to the control group might thus indicate a predisposition to a RISP episode. We also showed a slight nonsignificant increase in delta activity in REM and NREM 3. Interestingly, a similar finding, i.e. increased delta activity in patients compared to controls in REM and NREM 3 (uncorrected), was reported in a narcolepsy study [44]. 50-60 % of narcolepsy patients also report experiencing sleep paralysis episodes [45]. Further EEG differences in RISP patients’ REM sleep were described in our previous study [13]. Our current study shows that there are some unusual spectral power fluctuations in REM and also in NREM sleep. These findings support previous research [46, 47] about the relationship between sleep quality and the propensity to RISP episodes.

Next, EEG MS analysis was performed. Resulting topographical maps showed high GEV and visual similarity to canonical MS maps and we thus labeled them accordingly (A-D). The fact that the MS maps are coherent with maps found across studies [25] supports the power of this analysis. After the Bonferroni correction, we found lower transition rates between MS A-B and higher transition rates of MS C-D in RISP patients’ NREM 3 sleep stage. No significant differences were found in other temporal MS parameters (duration, occurrence, contribution, and GFP) and no other sleep stage was analyzed due to low correlations between template maps and individual subjects’ maps.

We’ve also shown an overall dominance of MS C and D topographies in RISP patients’ NREM sleep. MS A, B, and C were previously associated with sleep, but their temporal dynamics vary by states of vigilance and in psychiatric disorders [37]. Especially the duration parameter of MS is a frequent differentiating indicator of MS behavior [48]. We observed a slightly increased duration of MS C and MS D in RISP patients which may be pointing to a higher vigilance state in RISP patients. This duration difference, although not significant, may also be supported by the observed higher levels of theta activity during deep sleep. These results indicate the greatest stability of microstate C in terms of spatial arrangement. MS C duration parameters were previously also shown to be increased in narcolepsy patients during wake and reduced during sleep [45]. This study concluded that these microstructural EEG alterations might reflect the intrusion of brain states characteristic of wakefulness into sleep and instability of the sleep-regulating flip-flop mechanism, resulting not only in pathological switches between REM and NREM sleep but also within NREM sleep itself, which may lead to a microstructural fragmentation of the EEG [49].

In literature, MS C has the highest correlation across average maps in different datasets [20]. Transitions to MS C also often mark a pathological condition such as anxiety disorder, migraines, or dementia [50, 51, 52]. MS C was further associated with the cingulate cortex and cingulate gyrus [53]. The cingulate cortex was proposed to be the main mediator for switching between all MS classes [54, 55] as it plays an important role in switching between vigilance states [37].

According to a study by [56], MS C and D show the highest occurrences in NREM sleep data. Source localization showed the sources of MS C to be in frontal brain regions and sources of MS D in occipital areas. MS C may signal physiological frontal area deactivation typical for deep sleep. The increased number of transitions between our MS maps could reflect this switching mechanism between activation-inactivation states. Furthermore, increased transitions between MS associated with vigilance may contribute to a manifestation of a RISP episode. However, caution is in order when evaluating EEG-fMRI MS analyses due to possible artifact residuals caused by MRI which may be a contributor to the MS maps [57].

In terms of the strict association of sleep paralysis with REM sleep, it is also interesting that the association of another nearby REM parasomnia, as is nightmare disorder with REM sleep, which is a significant predictor of sleep paralysis [18], has recently become questionable. Literature states that there is no solid evidence indicating that nightmares occur more frequently during REM sleep than during NREM sleep. For instance, nightmares in PTSD patients are evenly distributed across NREM and REM periods [58]. There are several recent studies investigating neurophysiological correlates of nightmares, irrespective of the influence of comorbid psychopathologies [59]. Subjects who recall frequent nightmares were characterized by increased arousal-related cortical activity [60] and enhanced high-frequency electroencephalographic (EEG) oscillations during NREM sleep compared to control participants [16, 17]. The altered arousal-related activity was more pronounced during transitions between NREM and REM sleep states [61], suggesting that NREM-REM sleep transitions facilitating microarousals, microsleeps, and sleep interruptions may be specifically disrupted in these individuals. Interestingly, we also found altered microstate dynamics in NREM sleep, again suggesting a more complex sleep dysregulation beyond REM sleep in RISP patients. To validate our findings and further elucidate the RISP pathophysiology it would be convenient to measure subjects during sleep paralysis episodes and then compare the power changes in studied frequency bands and also focus on the characteristics of MS, all in the deep sleep phase. To conclude, our study shows that EEG abnormalities in RISP patients are present outside paralysis episodes and are not necessarily present only in REM sleep. Future studies could therefore further investigate these traces of RISP beyond REM sleep.

## Acknowledgements

This work was supported by the Grant Agency of the Czech Technical University in Prague, reg. No. SGS22/200/OHK4/3T/17 and reg. No. SGS21/140/OHK4/2T/17 and by programme Cooperatio Neurosciences of Charles University.

